# Evaluating image upsampling strategies for downstream microscopy image classification

**DOI:** 10.64898/2026.02.13.705844

**Authors:** Sakib Mohammad, Aalvee Asad Kausani, Md Noumil Tousif

## Abstract

Microscopy images are frequently downsampled due to acquisition and computational constraints, requiring reconstruction before downstream analysis. While super-resolution (SR) is typically assessed using pixel-level fidelity metrics, its impact on deep learning (DL) model behavior remains insufficiently understood. In this work, we present a study that examines how different upsampling strategies affect image quality and classification performance. Using the BloodMNIST dataset, we construct matched 224×224 datasets from 64×64 images via bicubic interpolation, SwinIR Classical, and SwinIR RealGAN DL SR models, alongside the original 224 ground-truth images. We evaluate reconstruction quality using the Structural Similarity Index Measure (SSIM) and Peak Signal-to-Noise Ratio (PSNR) scores and assess downstream classification performance using ResNet-50 and Vision Transformer models, with accuracy, macro-F1 score, and a confidence-aware metric, the area under the receiver operating curve for successful prediction (AUPR Success). Our results demonstrate that bicubic interpolation significantly degrades classification performance, whereas SR methods can recover class-relevant information, even better than the ground-truth data. These findings emphasize the importance of confidence-aware evaluation and unambiguous reporting of reconstruction pipelines in microscopy-based DL studies.

## I. Introduction

Deep learning (DL) models for microscopy are typically trained and evaluated at the resolution at which the images were acquired [1]. However, in practice, microscopy pipelines often involve changes in resolution, downsampling for storage and throughput, or upsampling for visualization. These transformations can alter fine-grained morphological features (texture, edges, grain) that are crucial for reliable classification of cell phenotype. As a result, a model’s apparent performance can reflect not only its representational capacity, but also the image formation pathway used to present input at a target resolution. In conjunction, we ask: How do common upsampling and DL-based super-resolution (SR) techniques affect pixel-level fidelity, downstream classification [2] performance, and prediction confidence? Addressing this fundamental question is important for comparative studies, as conclusions about classification performance depend on the data preprocessing method used to scale the input size.

In this work, we present a comparative study using the BloodMNIST dataset, a subset of the MedMNIST family of datasets [3] exported into four conditions at a common spatial size (224×224): (i) the original ground-truth 224 images (GT), (ii) bicubic upsampling (bicubic) to 224, (iii) SwinIR [4] Classical outputs, and (iv) SwinIR RealGAN [5] outputs. We evaluate these images from two perspectives:

i. Image fidelity using Structural Similarity Index Measure (SSIM) [6] and Peak Signal to Noise Ratio (PSNR) to quantify how closely the reconstructed images match the ground-truth.
ii. Downstream classification performance using two computer vision models, namely the ResNet-50 [7] and Vision Transformer Base with 16×16 patch size (ViT-B) [8], evaluated with accuracy, macro-F1 [9] and a confidence of correctness metric, namely area under receiver operating curve [10] for successful predictions (AUPR Success) [11] that treats correct prediction as the positive event and uses softmax score as confidence.

We note that we do not aim to maximize raw classification performance. Rather, we intentionally adopt a lightweight training regimen and strict seeding to observe differences in scores for the dataset image reconstruction methods only.

In the remainder of the paper, we describe the reconstruction pipeline and training protocol (Figure 1), then report the quantitative fidelity scores (Table 1) and downstream classification outcomes (Table 2), and finally analyze confidence shifts on a fixed set of test images (Figure 2) to connect reconstruction artifacts with prediction behavior.

**TABLE 1.**
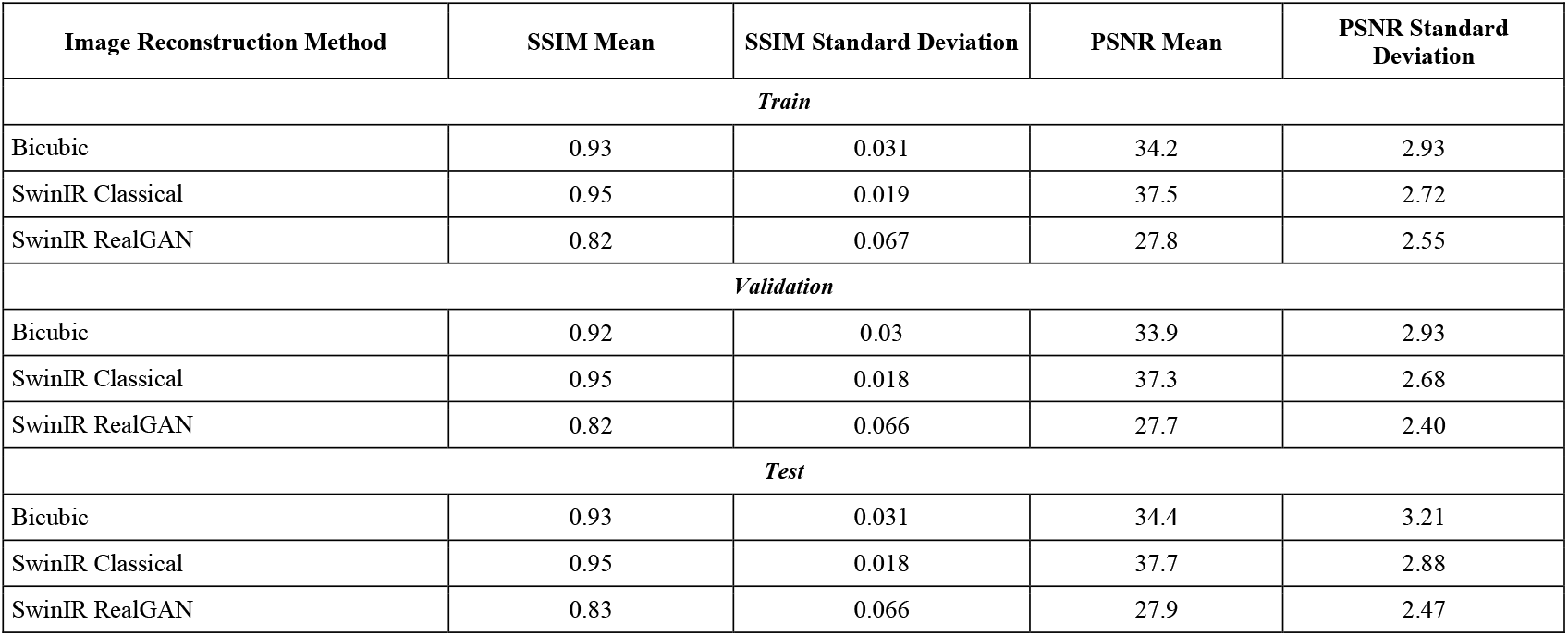
Image fidelity assessment scores across reconstruction methods and data splits.

| Image Reconstruction Method | SSIM Mean | SSIM Standard Deviation | PSNR Mean | PSNR Standard Deviation |
| --- | --- | --- | --- | --- |
| *Train* |  |  |  |  |
| Bicubic | 0.93 | 0.031 | 34.2 | 2.93 |
| SwinIR Classical | 0.95 | 0.019 | 37.5 | 2.72 |
| SwinIR RealGAN | 0.82 | 0.067 | 27.8 | 2.55 |
| *Validation* |  |  |  |  |
| Bicubic | 0.92 | 0.03 | 33.9 | 2.93 |
| SwinIR Classical | 0.95 | 0.018 | 37.3 | 2.68 |
| SwinIR RealGAN | 0.82 | 0.066 | 27.7 | 2.40 |
| *Test* |  |  |  |  |
| Bicubic | 0.93 | 0.031 | 34.4 | 3.21 |
| SwinIR Classical | 0.95 | 0.018 | 37.7 | 2.88 |
| SwinIR RealGAN | 0.83 | 0.066 | 27.9 | 2.47 |

**TABLE 2.** Classification performance of the dl models for dataset variants for the test set.

| Dataset Variant | Accuracy | F1 Score | AUPR Success |
| --- | --- | --- | --- |
| *ResNet-50* |  |  |  |
| Ground Truth | 0.9640 | 0.9650 | 0.9972 |
| Bicubic | 0.9623 | 0.9600 | 0.9972 |
| SwinIR Classical | 0.9664 | 0.9631 | 0.9977 |
| SwinIR RealGAN | 0.9743 | 0.9735 | 0.9977 |
| *ViT-B* |  |  |  |
| Ground Truth | 0.8901 | 0.8743 | 0.9819 |
| Bicubic | 0.8720 | 0.8548 | 0.9769 |
| SwinIR Classical | 0.8913 | 0.8766 | 0.9818 |
| SwinIR RealGAN | 0.8962 | 0.8831 | 0.9826 |

**Fig. 1.**
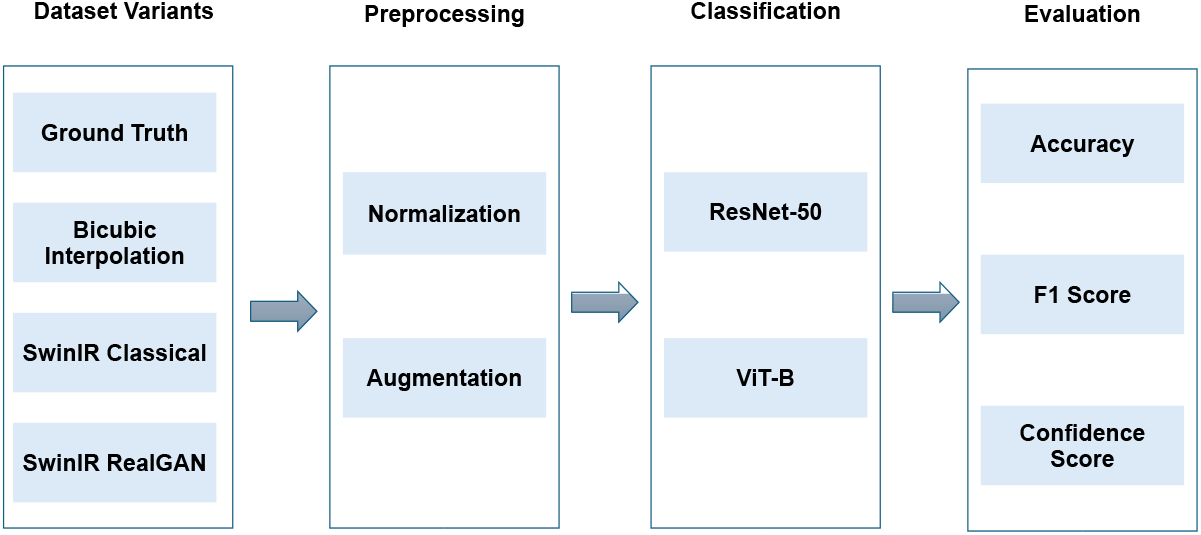
Overview of the experimental pipeline used to evaluate the impact of image reconstruction strategies on downstream microscopy image classification

**Fig. 2.**
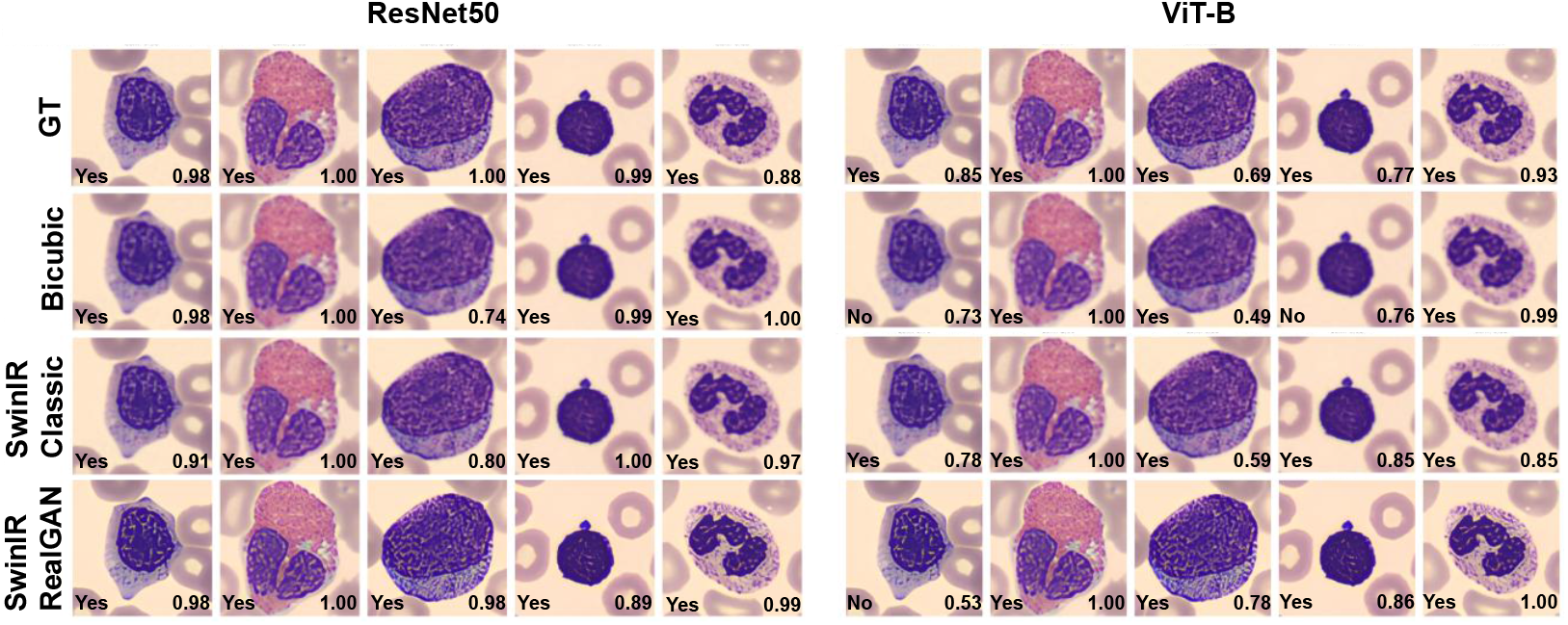
Effect of image reconstruction strategy on prediction outcome and confidence for representative test samples. Bottom left text on each image indicates whether it was correctly predicted, while the bottom right text specifies the confidence score

## II. Methods

### A. Dataset and Variants

All experiments were conducted on the BloodMNIST dataset, a multi-class microscopy benchmark consisting of RGB-images of peripheral blood cell types. To enable a fair comparison across image reconstruction methods, we constructed four dataset variants, all standardized to a spatial resolution of 224×224 pixels and sharing an identical train, validation, and test split:

(i) GT: Native 224×224 images provided by BloodMNIST.
(ii) Bicubic: Native 64×64 images upsampled to 224×224 using bicubic interpolation
(iii) SwinIR Classical: Images reconstructed from native 64×64 using a SwinIR Classical SR model and then resized to 224×224.
(iv) SwinIR RealGAN: Images reconstructed from native 64×64 using a SwinIR RealGAN-based SR model and then resized to 224×224.

Labels were exported directly from the official MedMNIST application programming interface (API) using the image index, ensuring consistency across dataset variants.

### B. Super-Resolution Models

We employed SwinIR, a transformer-based image restoration framework that leverages shifted-window self-attention [12] to capture both local and global dependencies. Two pretrained SwinIR models were used:

(i) Classical: Originally trained on paired low- and high-resolution images with a pixel-wise reconstruction objective, optimized for high SSIM and PSNR.
(ii) RealGAN: Trained originally using adversarial objective [13] and perceptual losses [14], designed to generate visually plausible and realistic textures rather than optimizing for fidelity scores.

The rationale for selecting these two models was to compare fidelity-oriented reconstruction against perceptual realism-inspired reconstruction. Both models performed ×4 upsampling from 64 to 256, after which images were resized to 224 for consistency across datasets.

### C. Image Fidelity Evaluation

To quantify reconstruction quality, we computed SSIM and PSNR between reconstructed images and the GT. Metrics were computed using standard formulations. For SSIM:

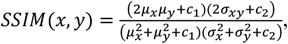

where *µ*_*x*_ is the pixel sample mean of *x, µ*_*y*_ is the pixel sample mean of *y*, 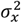 is the sample variance of *x*, 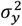 is the sample mean of *y, σ*_*xy*_ is the sample covariance of *x* and *y, c*_*1*_ = (*K*_1_*L*)^*2*^, *c*_2_ = (*K*_2_*L*)^*2*^ are two variables that stabilize the division with a weak denominator, *L* is the dynamic range of the pixel-values (typically calculated as 2^*no of bits per pixel*^ − 1), and *K*_1_ = 0.0*1, K*_*2*_ = 0.03 by default.

Finally, PSNR is given by,

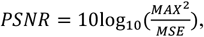

where *MAX* is 255 for an 8-bit image, and the mean squared error (MSE) is given by,

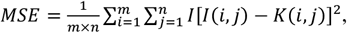

where *m, n* = dimensions of the image, and *I* is the original image, and *K* is the reconstructed image.

To reduce computational overhead while preserving statistical representation, metrics were estimated using random subsets of 200 images for the train, validation, and test splits, with a fixed random seed to ensure reproducibility.

### D. Classification Models

To study the downstream effect of reconstruction methods, we fine-tuned two image classification models: ResNet-50 and ViT-B. Both models were initialized with ImageNet [15] weights, and the final classification layer was replaced to match the classes of the BloodMNIST dataset.

### E. Training Protocol

We adopted a lightweight training strategy to avoid overfitting and emphasize comparative trends over peak performance. After initializing the models with ImageNet weights, we fine-tuned them with the Adam optimizer [16] with a learning rate of 1e-3 for 5 epochs. This choice was motivated by empirical observations that validation performance quickly saturated after 3-5 epochs. Extending training further would have harmed the comparison between the dataset variants. In addition, during training, we applied geometric image augmentations, including random horizontal and vertical flips (50% probability), random rotation (±15°), and random affine transformations (translation ±5%, scaling 0.95–1.05), to improve robustness while preserving cellular morphological features. Finally, all images were normalized using ImageNet mean and standard deviation.

### F. Evaluation Metrics

Classifier performance was evaluated on the held-out test set using accuracy and macro-averaged F1 score. To explicitly capture prediction confidence, we computed AUPR Success, defined as the area under the precision-recall curve where a prediction is labeled successful if it is correct, and the confidence score is the maximum softmax probability. Formally, if *s*_*i*_ ∈ {0,*1*} and *p*_*i*_ denotes confidence then,

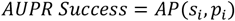

This metric assesses whether a model assigns high confidence to correct predictions, providing insights beyond accuracy alone.

Our experimental methods are summarized in Figure 1.

## III. Results and discussion

### A. Image Fidelity Assessment

Table 1 summarizes the pixel-level reconstruction quality of the three upsampling strategies relative to the GT images, using SSIM and PSNR across different dataset splits. We note that the SwinIR RealGAN model had the worst SSIM and PSNR scores, while the SwinIR Classical model had the best. It was expected, as the SwinIR Classical model was specifically trained to achieve high image fidelity, particularly in pixel-level reconstruction accuracy. On the other hand, the SwinIR RealGAN model produces the sharpest images among the strategies used here, at the expense of information. We observe a trade-off among SR models: perceptual and adversarial training in RealGAN favor realism at the cost of strict pixel fidelity. Finally, bicubic interpolation falls between the SR models, demonstrating its ability to recover high-frequency detail. The SSIM and PSNR metrics, while they quantify the closeness to actual GT data, cannot determine how a vision model will interpret the reconstructed images, motivating us for downstream evaluations beyond these metrics.

### B. Classification Performance

Table 2 reports downstream classification results for ResNet-50 and ViT-B across the dataset variants evaluated on the test set, using accuracy, macro-F1, and AUPR Success. We notice that ResNet-50 has better scores overall than ViT-B, as ResNet’s convolutional inductive biases help it learn visual features more quickly than ViT. With more training epochs, the ViT-B should reach the performance level of ResNet. Additionally, the SwinIR RealGAN dataset achieves the best scores across all metrics for both models, whereas the bicubic interpolation dataset yields the worst classification performance. Surprisingly, both SR datasets outperform the GT, indicating that reconstruction artifacts do not affect image classification. This finding is consistent with our previous work [17], in which using native 16-bit images (higher fidelity) rather than downscaled 8-bit images (lower fidelity) yielded no performance improvements in microscopy image classification on an in-house dataset.

From the results of the current experiments, we conclude that perceptually enhanced textures, even if less faithful pixel-wise, may improve model performance and confidence calibration. In addition, simple interpolation, while convenient, cannot preserve cellular textures and likely reduces discriminative cues, resulting in low performance.

### C. Per-Image Behavior

Figure 2 provides both a qualitative and a quantitative view of the prediction confidence for a fixed set of five test images, shared across all dataset conditions. Across most samples, we observed that confidence scores were the highest overall for the SwinIR RealGAN on ResNet-50, though it made one incorrect prediction on ViT-B (for the remaining 4 images, it had the highest overall confidence). Bicubic interpolation showed reduced confidence for ViT-B, even resulting in two incorrect predictions, suggesting that interpolation artifacts introduced uncertainty. For the SR variants, confidence patterns diverged in informative ways. SR Classical had the most robust performance among the image reconstruction methods, across the classification models (close or even better than GT for some samples), consistent with its high SSIM/PSNR.

This behavior further supports the notion that perceptual realism can positively influence the classifier’s internal representation and confidence estimation, even when pixel-wise similarity metrics are lower. Importantly, Figure 2 also reveals cases where confidence does not strictly track correctness across different methods. For example, despite having relatively high confidence scores (over 0.70) for samples 1 and 4 of the test images produced by bicubic interpolation, ViT-B made erroneous predictions. This reinforces the central dogma of this study: image reconstruction choices can systematically alter not only accuracy but also the confidence landscape of model predictions.

### D. Implications of Design Choices

Our results validate several design choices adopted in the study. The use of a short and fixed training regimen ensured that differences across dataset variants were not obscured by extensive fine-tuning. Similarly, strict seeding enabled fair comparisons across training conditions. The inclusion of the confidence criterion proved valuable as it revealed nuances that accuracy or F1 would have missed. We demonstrated that SR-based upscaling is not merely a preprocessing step but a factor that can meaningfully influence feature representation and learning. Evaluations that rely solely on pixel fidelity or classification accuracy risk overlooking important shifts in model confidence and behavior.

## IV. Conclusion

In this study, we conducted a controlled evaluation of how image upsampling strategies influence image fidelity and downstream behavior in microscopy image classification. By reconstructing four dataset variants from BloodMNIST, we ensured that all comparisons were performed under identical data splits, labels, model architectures, and training protocols.

Our results show that simple interpolation is insufficient to preserve class-relevant information. In contrast, DL-based SR reconstruction achieves superior performance, sometimes even better than the ground-truth images. We found that pixel-level fidelity is not sufficient to quantify downstream classification outcomes. Although SwinIR Classical achieved the highest SSIM and PSNR, SwinIR-RealGAN produced better overall classification results. These findings highlight that reconstruction pipelines can affect not only what a model predicts, but also how confidently it predicts, even when conventional accuracy metrics appear similar. For future work, we intend to refine the SR models by fine-tuning them on microscopic datasets before using them for upsampling. We also want to expand the number of classification models for a thorough evaluation. Finally, we wish to visualize the object localization of the classification models during inference to ensure they focus on relevant information in the images rather than artifacts. These additional experiments will help strengthen the claims made in this work.

## References

[1] Y. Zhou, J. Sollmann and J. Chen, “Deep-learning-based image compression for microscopy images: An empirical study,” Biological Imaging, p. e16, 2024.

[2] H. Su, Y. Li, Y. Fu and S. Liu, “A review of deep-learning-based superresolution: From methods to applications,” Pattern Recognition, vol. 157, 2025.

[3] J. Yang, R. Shi, D. Wei, Z. Liu, L. Zhao, B. Ke, H. Pfister and B. Ni, “MedMNIST v2 - A large-scale lightweight benchmark for 2D and 3D biomedical image classification,” Scientific Data, vol. 10, 2023.

[4] J. Liang, J. Cao, G. Sun, K. Zhang, L. V. Gool and R. Timofte, “SwinIR: Image Restoration Using Swin Transformer,” in CVF International Conference on Computer Vision Workshops (ICCVW), 2021.

[5] X. Wang, L. Xie, C. Dong and Y. Shan, “Real-ESRGAN: Training Real-World Blind Super-Resolution with Pure Synthetic Data,” Image and Video Processing, vol. 2, 2021.

[6] Z. Wang, A. C. Bovik, H. R. Sheikh and E. P. Simoncelli, “Image quality assessment: from error visibility to structural similarity,” IEEE Transactions on Image Processing, vol. 13, no. 4, pp. 600–612, 2004.

[7] K. He, X. Zhang, S. Ren and J. Sun, “Deep Residual Learning for Image Recognition,” Computer Vision and Pattern Recognition, 2015.

[8] A. Dosovitskiy, L. Beyer, A. Kolesnikov, D. Weissenborn, X. Zhai, T. Unterthiner, M. Dehghani, M. Minderer, G. Heigond, S. Gelly, J. Uszkoreit and N. Houlsby, “An Image is Worth 16×16 Words: Transformers for Image Recognition at Scale,” Computer Vision and Pattern Recognition, vol. 2, 2021.

[9] J. Opitz and S. Burst, “Macro F1 and Macro F1,” Machine Learning, vol. 3, 2019.

[10] R. J. Mark, B. Junge and J. R. Dettori, “ROC Solid: Receiver Operator Characteristic (ROC) Curves as a Foundation for Better Diagnostic Tests,” Sage Journals, vol. 8, no. 4, 2018.

[11] C. Corbière, N. Thome, A. Bar-Hen, M. Cord and P. Pérez, “Addressing Failure Prediction by Learning Model Confidence,” Computer Vision and Pattern Recognition, vol. 2, 2019.

[12] Z. Liu, Y. Lin, Y. Cao, H. Hu, Y. Wei, Z. Zhang, S. Lin and B. Guo, “Swin Transformer: Hierarchical Vision Transformer using Shifted Windows,” Computer Vision and Pattern Recognition, vol. 2, 2021.

[13] I. J. Goodfellow, J. Pouget-Abadie, M. Mirza, B. Xu, D. Warde-Farley, S. Ozair, A. Courville and Y. Bengio, “Generative Adversarial Networks,” Machine Learning, 2014.

[14] J. Johnson, A. Alahi and L. Fei-Fei, “Perceptual Losses for Real-Time Style Transfer and Super-Resolution,” Computer Vision and Pattern Recognition, 2016.

[15] O. Russakovsky, J. Deng, H. Su, J. Krause, S. Ma, S. Satheesh, A. Karpathy, A. Khosla, M. Bernstein, A. C. Berg and L. Fei-Fei, “ImageNet Large Scale Visual Recognition Challenge,” Computer Vision and Pattern Recognition, vol. 3, 2015.

[16] D. P. Kingma and J. L. Ba, “Adam: A method for stochastic optimization,” in Proc. Int. Conf. Learn. Represent. (ICLR), San Diego, CA, USA, 2015, pp. 1–13.

[17] S. Mohammad, “Deep learning powered identification and classification of early differentiated germ layers from pluripotent stem cells,” Ph.D. dissertation, Dept. of Electrical and Computer Engineering, Southern Illinois Univ. Carbondale, Carbondale, IL, USA, 2025. [Online]. Available: https://opensiuc.lib.siu.edu/dissertations/2345/

